# SeqWho: Reliable, rapid determination of sequence file identity using *k*-mer frequencies

**DOI:** 10.1101/2021.03.10.434827

**Authors:** Christopher Bennett, Micah Thornton, Chanhee Park, Gervaise Henry, Yun Zhang, Venkat S. Malladi, Daehwan Kim

**Author notes:** Email addresses: CB, MT, CP, GH, YZ, VM, DK.

## Abstract

With the vast improvements in sequencing technologies and increased number of protocols, sequencing is finding more applications to answer complex biological problems. Thus, the amount of publicly available sequencing data has tremendously increased in repositories such as SRA, EGA, and ENCODE. With any large online database, there is a critical need to accurately document study metadata, such as the source protocol and organism. In some cases, this metadata may not be systematically verified by the hosting sites and may result in a negative influence on future studies. Here we present SeqWho, a program designed to heuristically assess the quality of sequencing files and reliably classify the organism and protocol type. This is done in an alignment-free algorithm that leverages a Random Forest classifier to learn from native biases in *k*-mer frequencies and repeat sequence identities between different sequencing technologies and species. Here, we show that our method can accurately and rapidly distinguish between human and mouse, nine different sequencing technologies, and both together, 98.32%, 97.86%, and 96.38% of the time in high confidence calls respectively. This demonstrates that SeqWho is a powerful method for reliably checking the identity of the sequencing files used in any pipeline and illustrates the program’s ability to leverage *k*-mer biases.

## Introduction

Over the years, there has been an explosion in the applications of sequencing technologies and of sequenced organisms ^1,2^. Due to the variety, recent advancements, and the reduced costs of these technologies, there has been a substantial increase in the number of raw and processed read files produced ^3^. As the use of sequencing finds further applications, the depth of sequencing increases, and as more data becomes publicly available, proper storage, maintenance, and documentation become crucial. Fortunately, there are a number of public repositories where raw and/or processed files can be stored such as ENCODE and the Sequence Read Archive (SRA) ^4,5^. Some of these repositories are well maintained, requiring extensive validation of the submitted files, while others traditionally rely on user reporting. This has led to some inconsistencies and possible errors in experimental protocol and/or the species of origin in the metadata provided for some of the files ^6^. Indeed, It is well documented that errors propagating from these mislabeled calls in metadata do negatively impact data integrity ^7,8^. Furthermore, it is often important to ensure users have proper input files before running a time-intensive analysis, pipeline, or program. This need for a validation check extends to receiving data from less well-curated or private databases that may have less than ideal documentation.

To overcome these issues, some researchers have developed thresholds and other methods for filtering out files inconsistent with expected criteria or are otherwise suspicious when compared to the literature ^6,9^. However, these imposed restrictions limit the available data one can use for large analyses and are reliant upon the assumption that the criteria used to filter can catch all potentially erroneous files. The most accurate way of determining a file’s origin is to search through the originating source studies for indication of identity of the file. In fact, a previous study sought to validate files in these databases by word association in source texts with limited success ^9^. Some major limitations of these methods include their frequent inability to be applied to unpublished data and the excessive time-consumption during manual checking. An alternative method is to align the files to different species genomes starting with the reported species and ensure that the alignments match those expected from the experimental protocol. This method, while accurate, is computationally intensive and not as conducive on large scale data projects where thousands of files may be analyzed and thus thousands of alignments need to be performed.

Thus, we reasoned that a more rapid and resilient way to assess the identity of a sequence file is the use of sequences in an alignment-free algorithm. There have been a number of studies demonstrating the ability to leverage *k*-mer identity and frequency biases to distinguish species in metagenomics studies and to validate *de novo* genome assembly ^10,11^.

Here we present SeqWho, an accurate method for rapid validation of origin species and sequencing type from FASTQ(A) data and heuristically measuring basic read quality metrics. SeqWho exploits the principle of *k*-mer frequency biases between different genomes and regions of the genome in the differentiation of origin species and sequencing type ^12^. In this study, we demonstrate that SeqWho can accurately categorize the source species and technology from new sequencing data (greater than 95% on high confidence calls), using a Random Forest classification model.

Furthermore, SeqWho is designed to be a very rapid and efficient software, taking less than 30 seconds per file to run and having low memory requirements (approx. 750 MB). Taken together, we show that SeqWho is a powerful program that can reliably and quickly classify a diverse range of sequencing files for use in validation or downstream analysis preprocessing. SeqWho is open-source software freely available at https://github.com/DaehwanKimLab/seqwho.

## Results and Discussion

### Algorithmic Design

#### Model Selection and Measurement Determination

When designing the algorithm used by SeqWho, we first needed to determine classification-critical parameters such as the number of reads needed, the numeric determinants to be used, and the classification model. Previous studies in metagenomics and transcript qualification have demonstrated the ability to use frequency biases between *k*-mers as a method for making determinations ^13,14^. Thus, we started by calculating the frequencies of 1-7mers using only a portion of reads from the sequencing files. We used the smallest file sampling as possible to ensure rapid processing time. To this end, we determined that any selection beyond 25,000 reads produced diminishing returns for *k*-mer frequency array changes (Figure 1).

**Figure 1.**
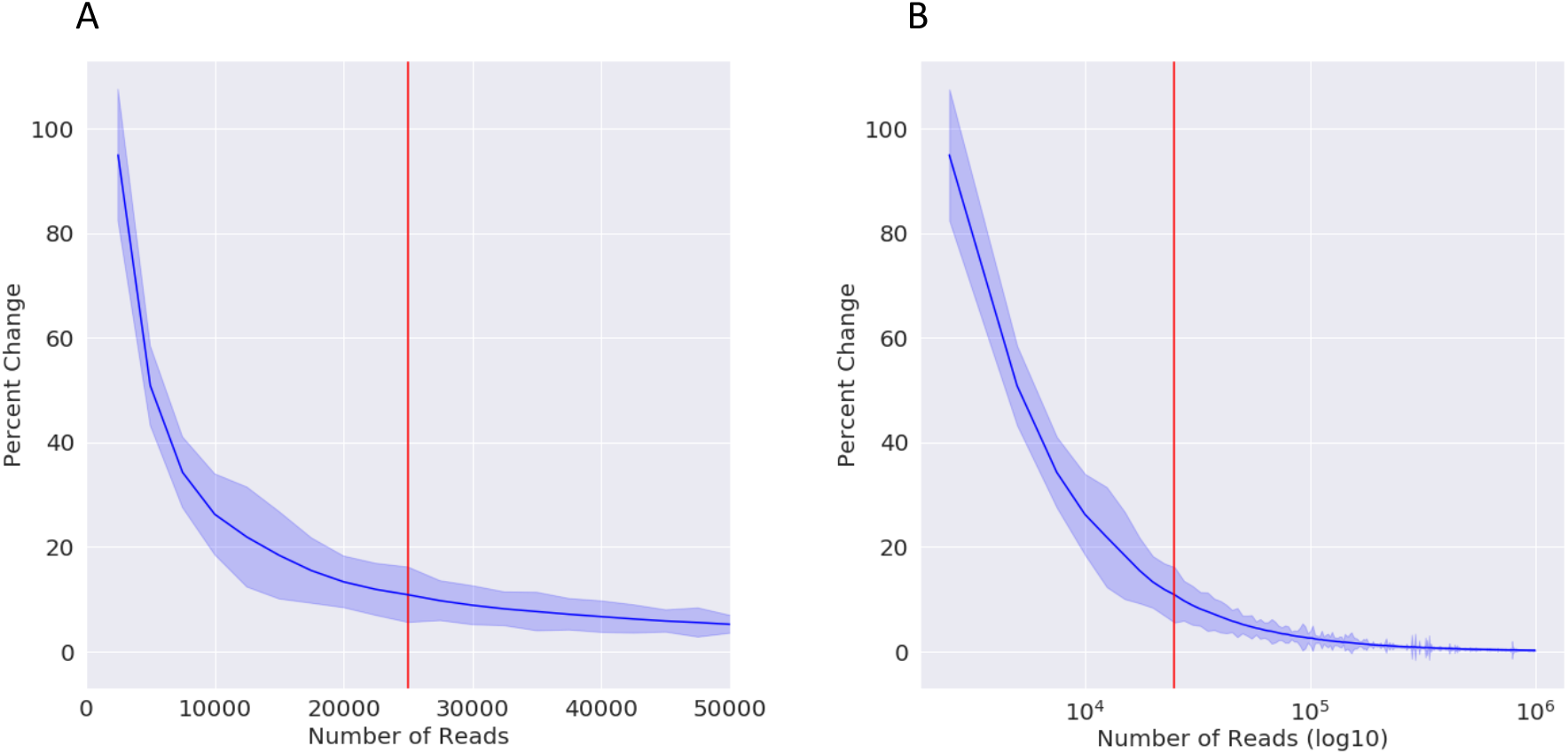
Plot of average percent change during k-mer table update as more reads are added. A) Shows number of reads maxed at 50,000 verses percent change. B) Shows log number of reads verses percent change. Red lines mark 25,000 reads

Next, we sought to test a number of supervised learning models on their ability to classify species of origin and sequencing type. We tested these models on 9 different sequencing technologies between two different species resulting in 18 total categories. We found that these initial 1-7mer frequency arrays were sufficient for better-than-random classification of species and sequencing type in all tested machine learning paradigms (Table 1). We chose to use the Random Forest Classifier as the workhorse of SeqWho’s algorithm due to its superior gains in classification accuracy with optimized parameters and ease of development.

**Table 1:** Classifier Performance. All classifiers tested with resulting call accuracy for both species and file type, species, and file type.

|  | Overall | Species | Type |
| --- | --- | --- | --- |
| **Random Forest** | 88.15% | 96.22% | 90.74% |
| **Neural Net Binary** | 84.80% | 95.58% | 87.64% |
| **Neural Net Categorical** | 84.41% | 95.85% | 87.34% |
| **Muli-Layer Perceptron** | 84.27% | 95.65% | 87.00% |
| **Logistic Regression** | 79.78% | 93.53% | 82.66% |
| **Linear SVM** | 70.21% | 84.31% | 76.79% |
| **k-Nearest Neighbor** | 70.15% | 93.15% | 77.69% |
| **Naïve Bayes** | 43.87% | 75.19% | 66.86% |
| **Decision Tree** | 37.28% | 85.22% | 63.64% |
| **Quadratic Discriminant** | 26.47% | 80.20% | 32.40% |

We then tested classification accuracies for various *k*-mer lengths to determine how many datapoints were needed for optimal classification (data not shown). While larger *k*-mers, around 31-mers, tended to produce better classification accuracy, the memory space needed to naïvely count *k*-mers at this level increases exponentially and makes the process very slow. Ultimately, we determined that 1-5mers were sufficient for classifying data at less than 90% overall accuracy. To improve this performance, we sought to include a small subset of common, highly deterministic 31-mers that are likely to appear within a sample of 25,000 reads of a FASTQ(A) file. Repetitive elements in the genome are very common and have been shown to be biased in species and genome location, and therefore useful in classification on species type and sequencing type ^15,16^. Furthermore, we recently developed HISAT2, a read alignment program that builds and utilizes a repeat element database from the genome, making this data very easy to obtain ^17^. Thus, we hypothesized that a combination of common *k*-mer indicators from repetitive genomic regions, designated here as repeat *k*-mers, as well as the initial frequency array would substantially enhance classification accuracy.

#### Resulting Design

We developed an algorithm to construct a training frequency array set using repeat 31-mers and 1-5 mers to train a set of core Random Forest Classifiers (Figure 2). We began with a set of FASTQ(A) files, labels, and HISAT2 repeat indices. For this initial test we used two different species, Human and Mouse, and nine different file types: Amplicon-seq, ATAC-seq, Bisulfite-seq, ChIP-seq, DNase-seq, miRNA-seq, RNA-seq, Whole Genome Sequencing (WGS), and Whole Exome Sequencing (WES).

**Figure 2.**
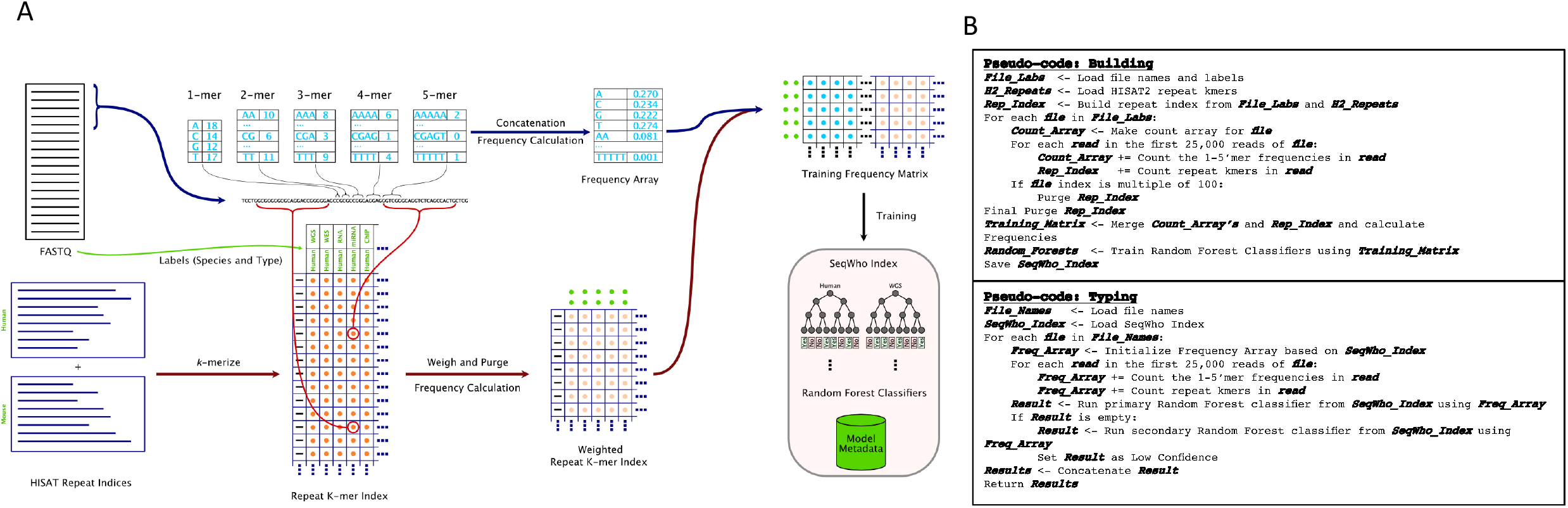
Model design for SeqWho Random Forest classification training. A) Graphical flow of data read in and processed into different arrays B) Pseudo-code for the whole process of building the indices needed and typing from the indices.

We attempted to download 1000 random files of each type from the SRA marked for public use totaling 18,000 files. However, some of the files failed to download, were outdated color-space reads, or had other formatting issues that lead to a loss of file integrity, mostly in the Human Amplicon category. We ended up with 17,489 total files of which 1,004 did not meet quality standards resulting in 16,485 files used in model training (Table 2).

**Table 2:** Number of files used in database training. Count of files of each species and file type used to build the original SeqWho model

|  | Amplicon<br>-seq | ATAC<br>-seq | Bisulfite<br>-seq | ChIP<br>-seq | DNase<br>-seq | miRNA<br>-seq | RNA-seq | Whole<br>Genome<br>Seq | Whole<br>Exome<br>Seq |
| --- | --- | --- | --- | --- | --- | --- | --- | --- | --- |
| Human | 621 | 1000 | 991 | 990 | 995 | 993 | 988 | 994 | 993 |
| Mouse | 1000 | 999 | 985 | 982 | 960 | 1000 | 1000 | 1000 | 998 |

We built 1-5mer frequency arrays and 31-mer repeat frequency arrays for each file and added them to a training frequency matrix (Figure 2A). To minimize the space needed to store repeat *k*-mers, after every 100 files processed, we purged the repeat *k*-mer index for any repeats that had less than a sliding threshold of hits. This allowed us to reduce the size of the index during building and limit repeats to only the most abundant in each file type. To keep from biasing the *k*-mer index purge to one type of file such as biasing to WGS, we randomized the selection of the files from the 16,485 pool. The resulting index contained 1095 of the most common repeat *k*-mers. These repeats were sorted and mapped to an array and frequencies were appended to the 1-5mer arrays. Through our multiple rounds of testing, we found that using binary Random Forest classifiers for each category were more accurate than categorical classification, easily reaching above 90% accuracy. However, we noted that there were some rare instances where no models were able to classify files and the inclusion of the combined classifications for species and type is necessary to serve as a second phase fail-safe for files that failed to be properly classified. Thus, the resulting frequency matrix was used to build 13 different Random Forest models, one for each classification: mouse, human, Amplicon, ATAC, Bisulfite, ChIP, DNase, microRNA, RNA, WGS, and WES; and two for categorical classifications, species and type. We also included metadata regarding model building so that the same steps can be used when typing incoming files against the index.

### Classification Results

We validated the model, trained using the aforementioned dataset, using 1,665 novel files (~100 of each species and sequencing type) not used during training. We found that we could correctly classify the species of the file ~98% of the time, sequencing technology ~95% of the time, and both combined ~93% of the time (Figure 3 and Table 3).

**Figure 3.**
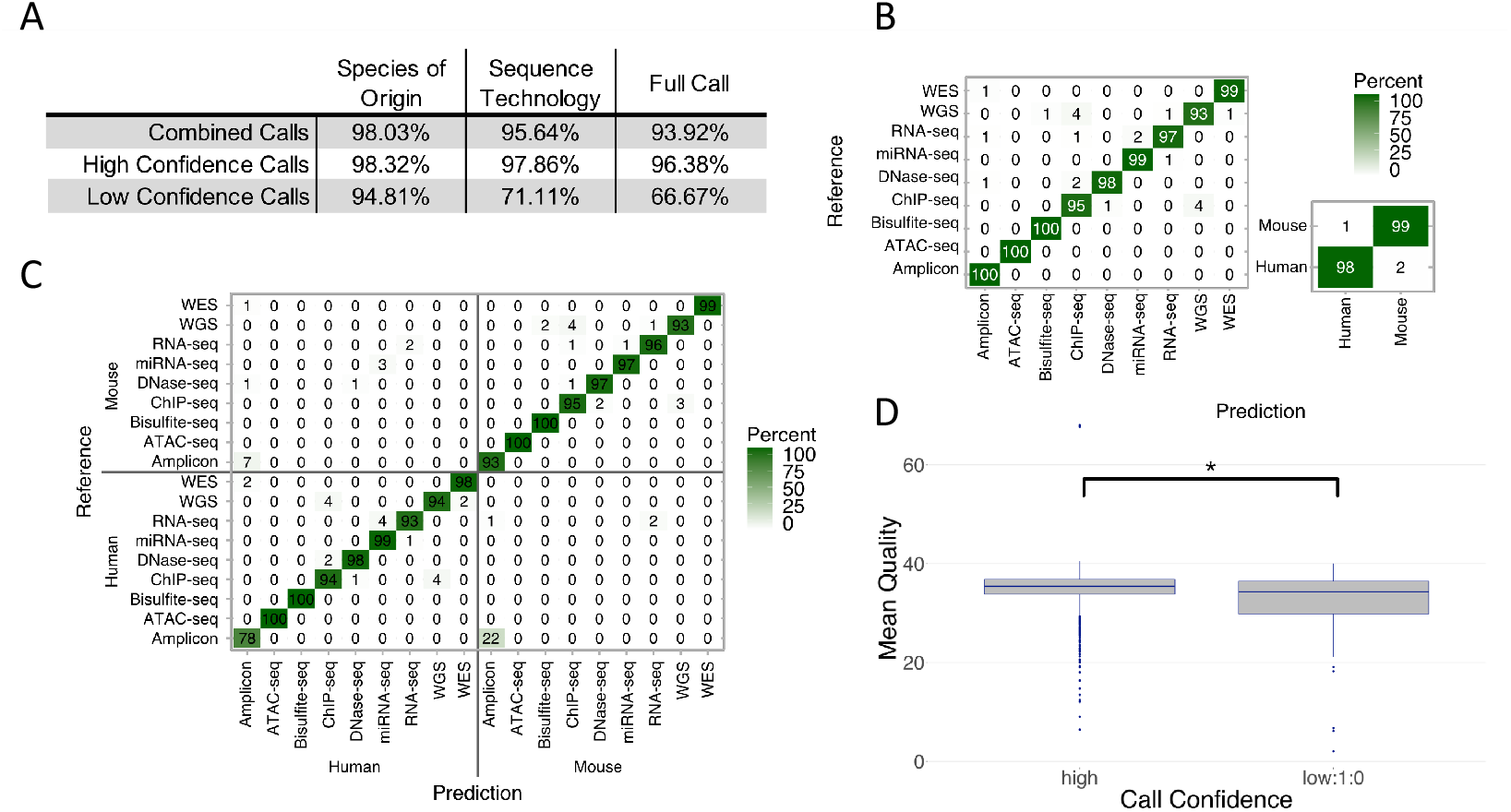
Classification accuracy of SeqWho’s Random Forest models. A) Percent correct calls for Species, Sequencing Type, and all together for all calls, high confidence calls, and low confidence calls. B) Confusion matrix for Species calls and Sequence type calls for high confidence calls in percent. C) Confusion matrix for full correct calls for high confidence calls in percent. D) Box-n-whisker plot showing differences between Mean Read Quality between high and low confidence calls. * indicates adjusted p < 0.0001

**Table 3:** Random Forest classification metrics. Metrics for random forest model in SeqWho. Equations for each metric are located in the column heading. TP = True positive, T = Total, TN = True Negative, N = Called Negative, P = Called Positive. Models were build using N=16,485 files (Table 2)

|  |  | Sensitivity/<br>Recall (S)<br>TP/T | Specificity<br>TN/N | Pos Pred Value/<br>Precision (PPV)<br>TP/P | Neg Pred Value<br>TN/N | F1<br>2SxPPV/(S+PPV) | Detection<br>Rate | Detection<br>Prevalence | Balanced<br>Accuracy |
| --- | --- | --- | --- | --- | --- | --- | --- | --- | --- |
| Human | Amplicon | 0.784 | 0.996 | 0.870 | 0.992 | 0.825 | 0.027 | 0.031 | 0.890 |
|  | ATAC | 1.000 | 1.000 | 1.000 | 1.000 | 1.000 | 0.064 | 0.064 | 1.000 |
|  | Bisulfite | 1.000 | 1.000 | 1.000 | 1.000 | 1.000 | 0.062 | 0.062 | 1.000 |
|  | ChIP | 0.943 | 0.996 | 0.930 | 0.997 | 0.936 | 0.044 | 0.048 | 0.970 |
|  | DNase | 0.976 | 0.999 | 0.976 | 0.999 | 0.976 | 0.054 | 0.055 | 0.987 |
|  | miRNA | 0.990 | 0.996 | 0.942 | 0.999 | 0.965 | 0.065 | 0.069 | 0.993 |
|  | RNA | 0.926 | 0.998 | 0.962 | 0.996 | 0.943 | 0.050 | 0.052 | 0.962 |
|  | WGS | 0.938 | 0.998 | 0.962 | 0.996 | 0.950 | 0.051 | 0.053 | 0.968 |
|  | WES | 0.985 | 0.999 | 0.970 | 0.999 | 0.977 | 0.043 | 0.044 | 0.992 |
| Mouse | Amplicon | 0.935 | 0.992 | 0.782 | 0.998 | 0.851 | 0.029 | 0.037 | 0.963 |
|  | ATAC | 1.000 | 1.000 | 1.000 | 1.000 | 1.000 | 0.066 | 0.066 | 1.000 |
|  | Bisulfite | 1.000 | 0.999 | 0.980 | 1.000 | 0.990 | 0.065 | 0.066 | 0.999 |
|  | ChIP | 0.955 | 0.996 | 0.913 | 0.998 | 0.933 | 0.042 | 0.046 | 0.975 |
|  | DNase | 0.967 | 0.999 | 0.989 | 0.998 | 0.978 | 0.058 | 0.059 | 0.983 |
|  | miRNA | 0.968 | 0.999 | 0.989 | 0.998 | 0.979 | 0.062 | 0.062 | 0.984 |
|  | RNA | 0.957 | 0.998 | 0.967 | 0.997 | 0.962 | 0.059 | 0.061 | 0.977 |
|  | WGS | 0.926 | 0.999 | 0.978 | 0.995 | 0.951 | 0.058 | 0.060 | 0.962 |
|  | WES | 0.990 | 1.000 | 1.000 | 0.999 | 0.995 | 0.064 | 0.064 | 0.995 |

Through this test, we observed that we could tag each call with a confidence based on which Random Forest models were used to make the classification. For example, files that only needed the first 11 binary classification models for a proper full classification producing a single species classification and a single technology classification were considered high confidence calls. In cases where classifications were absent, the categorical classification model was used and considered low confidence. Out of the 1,665 files, only two showed double sequencing type classification, and both cases were assigned a dual WGS and ChIP-seq classification with the truth set indicating WGS. This is not surprising as WGS files and ChIP-seq types are among the most commonly confused classifications in the model (Figure 3B-C). We suspect this is due to the *k*-mer and repeat bias in the intergenic regions of the genome being captured in both sets and used to determine classification. We found that the most difficult classification is human amplicon vs mouse amplicon with ~22% misclassified (Figure 3C). We are not surprised by this result as amplified regions can be highly diverse between different experiments and may lead to extraneous biases that confound the model’s ability to properly classify the files. Interestingly, we noted that RNA-seq also has some ambiguity in its classification of species. This could be due to the conservation of critical coding sequences between human and mouse genes. Overall, our Random Forest classifiers produce excellent results with high Sensitivity, Specificity, Precision, and Recall with the exception of Human Amplicon as mentioned above (Table 3).

To begin to understand what features determine whether a file will be classified high confidence or low confidence, we gathered read and file quality metrics and performed a student’s t-test between all numeric variables measured with a Bonferroni correction. We found that Mean Read Quality was significantly different between the two categories and may help explain why some files are easier to classify (Figure 3D). This result makes sense as lower quality reads may impact the *k*-mer bias between files since there is no error correction or *k*-mer omission in the model.

### Mixed Samples and Integration into Consortia RNA-seq pipeline

We wanted to stress-test SeqWho’s call function on data other than SRA data and apply SeqWho to a real-world use case for further development. An RNA-seq analysis pipeline for two NIH consortia was being developed and released during the same timeframe as SeqWho. This initiative experienced longer than ideal file validation due to using alignment-based validation methods. We partnered with the project lead to incorporate SeqWho and its prebuilt models into their process. We were able to confirm SeqWho’s processing time of approximately 20 seconds, 20-200 times faster than their previous method, and accuracy of greater than 95% confidence.

A regular challenge presented to the pipeline is of mixed or contaminated data (ie mouse data in human data). We tested SeqWho on a set of synthetic and real mouse, human, RNA-seq, and ChIP-seq data mixtures randomly generated or sampled from real data (Table 4). We found that in all cases where human sequences were present in mouse sequences, the data was classified as human. Furthermore, when RNA-seq data was present with ChIP-seq data, the call was always presented as RNA-seq. Interestingly, only a few of these calls had low confidence, indicating that the model may preferentially look for Human and DNA specific sequences or signatures over Mouse or RNA signatures. In one sense, this is surprising as we would expect an equal chance of Mouse or Human in a mixture. On the other hand, it is not surprising for the sequencing type call since RNA sequences are a subset of DNA sequences. It makes sense to look for DNA-specific markers. Of particular interest is the inclusion of single cell RNA-seq (scRNA) files in the test. Even though SeqWho was not trained on scRNA-seq files, it was able to accurately call the files as RNA-seq with one file having a low confidence tag.

**Table 4:** Mixed data type stress test. Table shows results of stress test on SeqWho using files containing mixed data. Mouse and Human RNA-seq and ChIP-seq used. Human:Mouse ratio is percent of file with human data with remainder representing mouse data. RNA-seq:ChIP-seq ration is percent of file with RNA-seq data with remainder representing ChIP-seq data. Single Cell RNA-seq indicates whether the sample is from single cell RNA-seq experiment. Call confidence is reported by SeqWho up arrow = high confidence, down arrow = low confidence. Call species and call sequence type are the results from SeqWho. Count is number of iterations that result was found in the combination of the previous columns of the table.

| Human :<br>Mouse<br>Ratio | RNA-seq :<br>ChIP-seq<br>Ratio | Single Cell<br>RNA-seq | Call<br>Confidence | Call Species | Call<br>Sequence<br>Type | Count |
| --- | --- | --- | --- | --- | --- | --- |
| 1.0 | 0.0 | ✗ | ↑ | Human | <u>ChIP-seq</u> | 1 |
| 1.0 | 0.3 | ✗ | ↓ | Human | RNA-seq | 5 |
| 1.0 | 0.4 | ✗ | ↓ | Human | RNA-seq | 5 |
| 1.0 | 0.5 | ✗ | ↓ | Human | RNA-seq | 2 |
| 1.0 | 0.5 | ✗ | ↑ | Human | RNA-seq | 3 |
| 1.0 | 0.6 | ✗ | ↑ | Human | RNA-seq | 5 |
| 1.0 | 0.7 | ✗ | ↑ | Human | RNA-seq | 5 |
| 1.0 | 1.0 | ✗ | ↑ | Human | RNA-seq | 2 |
| 0.7 | 0.3 | ✗ | ↑ | Human | RNA-seq | 1 |
| 0.7 | 0.3 | ✗ | ↓ | Human | RNA-seq | 4 |
| 0.7 | 1.0 | ✗ | ↑ | Human | RNA-seq | 5 |
| 0.6 | 0.4 | ✗ | ↑ | Human | RNA-seq | 5 |
| 0.6 | 1.0 | ✗ | ↑ | Human | RNA-seq | 5 |
| 0.5 | 0.5 | ✗ | ↑ | Human | RNA-seq | 5 |
| 0.5 | 1.0 | ✗ | ↑ | Human | RNA-seq | 5 |
| 0.4 | 0.6 | ✗ | ↑ | Human | RNA-seq | 5 |
| 0.4 | 1.0 | ✗ | ↑ | Human | RNA-seq | 5 |
| 0.3 | 0.7 | ✗ | ↑ | Human | RNA-seq | 5 |
| 0.3 | 1.0 | ✗ | ↑ | Human | RNA-seq | 5 |
| 0.0 | 1.0 | ✗ | ↑ | <u>Mouse</u> | RNA-seq | 11 |
| 0.0 | 1.0 | ✗ | ↑ | <u>Mouse</u> | RNA-seq | 11 |
| 0.0 | 1.0 | ✓ | ↓ | <u>Mouse</u> | RNA-seq | 1 |
| 0.0 | 1.0 | ✓ | ↑ | <u>Mouse</u> | RNA-seq | 1 |

### Heuristic FASTQ(A) quality information

Our goal is to make SeqWho as useful as possible for upstream processes in all major bioinformatic pipelines. Thus, we wanted to further expand SeqWho by reporting quality metrics on the files it processes, modeling the reports of another popular QC program FASTQC ^18^. The main metrics we focused on were: 1) %GC content 2) average read quality 3) number of reads in the file and 4) sequence/adapter content (Figure 4). These metrics were easily added to the processing steps taken when constructing frequency vectors with constant time changes to the algorithm. We wanted to avoid processing all reads in the file and maintain a rapid processing time for SeqWho’s as opposed to FASTQC which processes all reads in the file and is many times slower. However, while 25,000 reads are sufficient for classification, we wanted to make sure we captured sufficient quality information resulting in doubling the read number to 50,000. Compared to FASTQC as run on the 17 Platinum Whole Genome Sequences previously reported ^19^, we found that SeqWho is ~200 times faster and has very similar percent GC and average read quality metrics (Figure 4A). Though we added a naïve adapter detection step, SeqWho was not able to detect any adapters in the files tested. This may be a byproduct of the more stringent cutoffs SeqWho uses to assess bias in the ends of the reads, or a byproduct of focusing on the end 10 nucleotides of the reads for detection.

**Figure 4.**
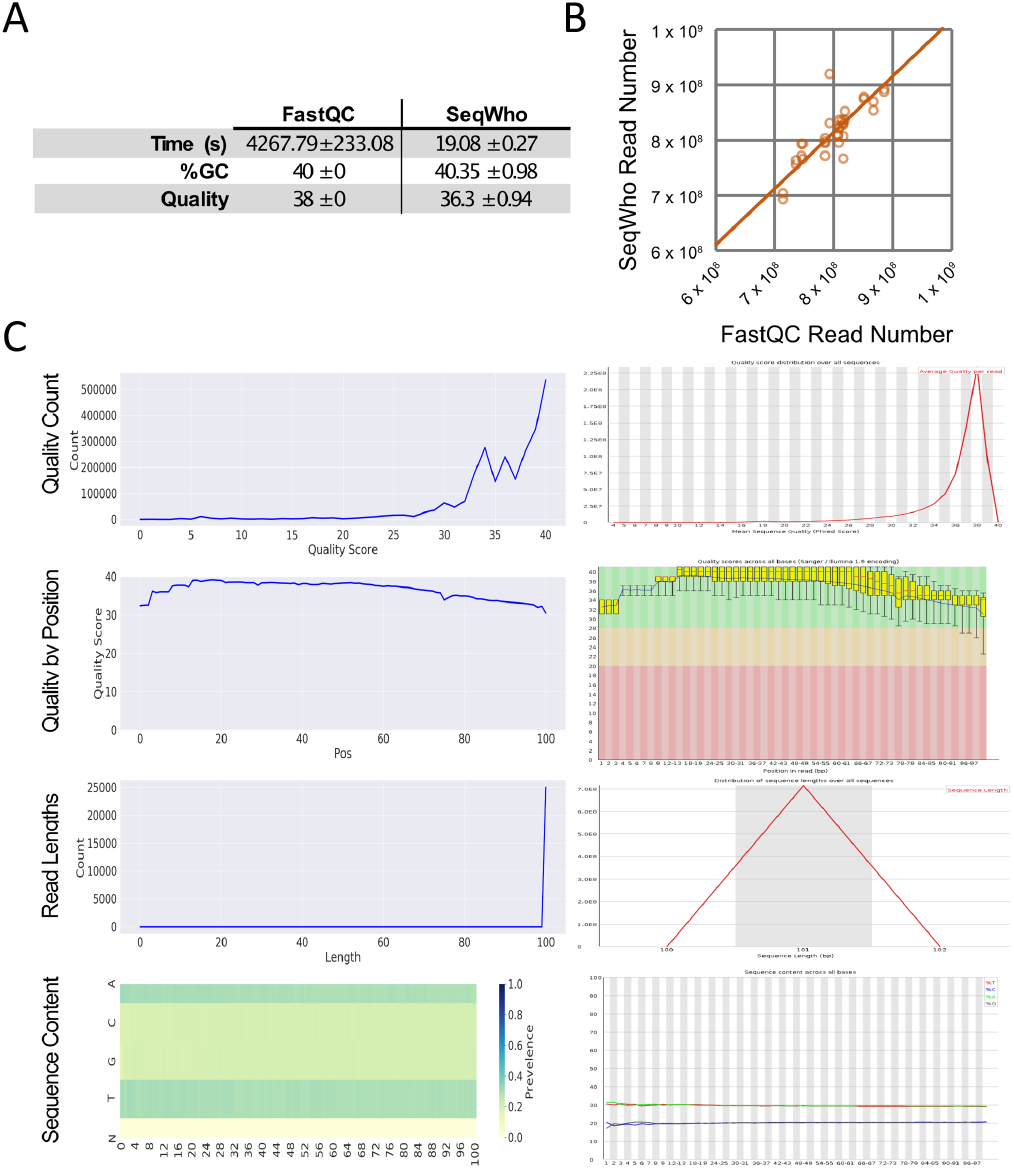
Comparison of SeqWho quality information to FASTQC. A) Comparison of times, %GC, and average read quality with standard deviations. B) Linear correlation between the true FASTQC file read number and the estimated SeqWho read numbers. C) Plot and data distribution comparison between SeqWho (left) and FASTQC (right).

We added a heuristic estimation of number of reads in a file using the ratio of the number of reads in a chunk to the size of the chunk and the size of the file. Interestingly, this method produces read estimates very similar to the true value as captured by FASTQC (Figure 4b). Only one file’s estimates did not coincide with the true read numbers. We suspect this was due to less efficient compression of a part of the file that threw off the ratio of our estimate.

Furthermore, we added plots to represent the quality information in a manner similar to FASTQC (Figure 4C). All plots that SeqWho produces contain highly similar data and trends as those produced by FASTQC. Thus, SeqWho can rapidly and reliably capture representative quality metadata from processed files. Additionally, this information can be read visually or by a computer program as all raw data is exported as a JSON and TSV file for later use.

## Conclusions

Here we presented SeqWho, a rapid and reliable software for classifying a read file’s original organism and sequencing type, and for assessing quality information. We utilized the bias in *k*-mer frequencies to train 13 Random Forest classifier models. This provides us with a reliable way to assess the confidence of the call and allows us to achieve upwards of ~97% accuracy in high confidence classifications. Furthermore, SeqWho allows us to rapidly assess the quality metrics of the reads and file as a whole with constant time addition to the algorithms. By using only 25,000-50,000 reads, we were able to keep the run time of SeqWho to ~20 seconds, ~200 times faster than another commonly used QC program, FASTQC, with the additional ability to classify the file. Additionally, SeqWho runtime is independent of input file size due to subsampling. While there are some errors in the heuristic assessment of quality, SeqWho remains able to very accurately characterize the file’s quality substantially faster than FASTQC. Furthermore, we report this data in a graphical format for human interpretation and as a JSON-formatted text file to be read in downstream automated processes. We consider this aspect to make SeqWho a critical and versatile program for use in standard sequence QC, in large scale data pipelines with extensive automation, or in individual cases to confirm data from dubious origins. Future work will focus on improving the algorithms classification to achieving the desired goal of >99% accuracy as well as improving the heuristic determination of quality information. Further implications of this work include that we are capable of drawing highly valuable information can be drawn from biases in *k*-mer frequencies without the time expensive step of read alignment. Overall, SeqWho is a versatile, rapid, and reliable program that lays the framework for extensive future work into utilizing *k*-mer frequency and repeat information in unique, rapid ways.

## Materials and Methods

### Program versions and code development

Unless otherwise noted, SeqWho was developed in a conda environment under python version 3.7.4 using package versions noted in supplemental_file_1.txt file. All codes, analyses, and plots were performed or developed within this environment on a Linux workstation running CentOS version 7 with a Xeon^®^ E5-1650 3.60 GHz 12 core CPU and 64GB of ECC RAM. In addition to python, we used Nextflow version 0.31.0, SRA-toolkit version 2.10.9, seqtk version 1.3, R version 4.0.3, FASTQC version 0.11.8, Keras version 2.2.4, and tenserflow version 1.14.0.

### Design Principles

#### Model Building

Model building is divided into two minor processes: 1) the 1-5 mer frequency generation and 2) repeat index consolidation. A list of training files is first read from a directory specified by the user and randomized. Then for each file for each read in the first 25,000 reads every 1-, 2-, 3-, 4-, and 5-mer without ambiguous nucleotides are counted and each count is added to a 1,364 long array position corresponding to a sorted list of *k*-mers. The resulting count array is converted to *k*-mer frequencies by *k*-mer set size and added to a matrix with the file label recorded. The second process, repeat index consolidation, involves building a sorted array of repeat 31-mers from the repeat indeces for mouse and human provided by HISAT2 v2.2. As each file is processed, each read is scanned for 31-mers matching sequences in the repeat array and counted in a count array. After every 100 processed files any repeat with a total number of hits less than the number of files processed divided by 100 are removed and their corresponding entries are removed from all other file count arrays. After all files have been processed a final purging step is performed to remove repeats with high similarity between species using the variance of the repeat frequencies. Any variance less than half the variance of a perfectly determining repeat (frequency of 1 for a single species) is removed. The resulting counts were converted into a frequency by dividing the counts for each file by the sum of the file counts and were added to the 1-5 mer frequency matrix. From this model building data, a binary Random Forest classifier was trained for each label and species, resulting in our case with 9 sequencing type and 2 species models. Two further models were trained using all sequencing type classification and all species classification resulting in a total of 13 Random Forest models with the result from each mapped to a result array. Metadata, including information needed to rebuild the vectors and repeat information, and the Random Forest classifiers in a python pickle were saved into a SeqWho index for use in file testing.

#### File Testing

File classification makes use of the metadata present from the building step to assemble a compatible frequency array that can be used in the Random Forest classifiers using steps identical to the building step except for the repeat purging procedure, which is not needed. Furthermore, read quality metrics: length, quality per base, average quality, nucleotide biases etc. were measured to be reported simultaneously to *k*-mer counting. An estimate of the total read number was calculated by multiplying the total number of reads read processed by the ratio between the total compressed size of the file on disk and the compressed size of the file chunk read in by Seq-Who. The binary classifiers were used first to determine if an accurate call can be made. If 1 species and 1 sequencing type call were generated the quality metrics and call were reported. Any other call generated results in a double check against the backup categorical Random Forest classifiers for species and sequencing type. In this case a low confidence flag was appended and included tag to delimitate which classification (species or sequencing type) had to be validated. A number one indicated that classification was a high confidence validation and a zero indicatred a lower confidence classification with the first number indicating species classification and the second number indicating sequencing type classification (ex “low confidence1:0” indicated species was called with high confidence and sequencing type needed to be validated). Calls and quality data are returned as a JSON and tab delimited text file with read quality information also being reported in a plot PNG file.

### Acquisition of Data

A Nextflow script was used to download specific sequencing read files from the SRA database. A metadata file for SRA, obtained from https://ftp.ncbi.nlm.nih.gov/sra/reports/Metadata/, was filtered for publicly available data. 19,800 files corresponding to whole genome sequencing, whole exome sequencing, ChIP-seq, amplicon sequencing, ATAC-seq, DNase-seq, Bisulfite-seq, RNA-seq, and micro-RNA-seq for each of two species, Mouse and Human, 1,100 files for each type, were randomly selected for download. We downloaded 1 million reads from each selected file using SRA-toolkit with the –maxSpotId option and Nextflow. The file retrieval process was run using the BioHPC (UT Southwestern) and up to ten files were retrieved simultaneously. The final database consisted of a total of 18,151 FASTQ(A) files gzip compressed, consuming approximately 164 GB of disk space. Data for the RNA-seq pipeline validation can be found at https://doi.org/10.5281/zenodo.4429315.

### Model Testing and Validation

Model testing and selection was performed using scikit-learn for the following models: Logistic Regression, Decision Tree, Naïve Bayes, k-Nearest Neighbor, Linear SVM, Multi-Layer Perceptron, Quadratic Discriminant, and Random Forest. Keras with Tenserflow was used for building and testing Neural Nets. To rapidly test the models and parameters, 100 files for each file sequencing type for each species were used to build a pre-calculated 1-7-mer count matrix with labels. To determine the number of reads needed, reads were drawn from the first 1 million reads in 100 increments and counts were added to the matrix. At each increment, frequencies were calculated by dividing the individual counts by the total counts and the percent change was calculated between the previous increment and the current increment. 25,000 reads were selected as sufficient for model testing. Thus, the count matrix was built with the top 25,000 reads of each file. Two frequency matrices were generated from the count matrix, with one, calculated by dividing the individual counts by the total counts and the other calculated by dividing each *k*-mer count set by the sum of the set. Each model was tested multiple times using an 80/20 split of each matrix with varying optimization parameters, levels, nodes, etc. RandomForestClassifier module was selected for use with n_estimator parameter set to 500.

The complete SeqWho algorithm was tested by building an index from the aforementioned SRA data. 16,485 files (Table 2) were used to build the model while the remaining 1,665 files were used for validation. Confusion matrices were built in R using the caret package version 6.0 and plotted using ggplot2 version 3.3.3. Statistics performed were student’s t-test with Bonferroni correction to account for multivariate testing. Seq-Who quality metrics were tested against FASTQC on 17 Platinum Genome sequencing files that we reported on previously ^17^.

Four replicate’s FASTQ’s were manually downloaded from the GUDMAP consortium data-hub website. The replicates represent three different sequencing modalities (bulk RNA-seq, scRNA-seq, and ChIP-seq), as well as two species (human, and mouse). To create RNA-seq (bulk) and ChIP-seq as well as - human and mouse admixtures, the FASTQ’s were randomly sampled and concatenated in order to generate varying amounts of sequence type and species mixtures. Each mixture was then randomly sampled to one million reads, using different seeds to create multiple replicates of the same admixtures. Seqtk (version 1.3) was use for the random sampling of the FASTQ’s. The resulting FASTQ’s were then analyzed using SeqWho to call sequencing types and species.

## Acknowledgements

This work was supported in part by the National Institute of General Medical Sciences (NIH) under grants R01-GM135341 and by the Cancer Prevention Research Institute of Texas (CPRIT) under grant RR170068 to D.K. Work for the RNA-seq pipeline was supported by NIH 5U24DK110814-05 and CPRIT grant Cancer Prevention and Research Institute of Texas (RP150596). All authors read and approved the final manuscript.

## Authors’ contributions

C.B., M.T., C.P., G.H., Y.Z, V.M., and D.K. performed validation analysis and discussed the results of SeqWho. C.B., and D.K. designed and implemented Seq-Who. G.H., and V.M. integrated SeqWho into RNA sequencing analysis pipeline and performed stress tests. C.B., M.T., Y.Z., and D.K. wrote the manuscript.

## Competing interests

The authors declare no competing financial interests.

## Availability of data and materials

Project name: SeqWho

Project home page: https://daehwankimlab.github.io/seqwho

Operating system(s): Linux, Mac OS X and Windows

Programming language: Python

License: GPLv3 license

## Notes

https://github.com/DaehwanKimLab/seqwho

### Competing Interest Statement

The authors have declared no competing interest.

